# DNA methylation in *Escherichia coli* changes in response to growth conditions

**DOI:** 10.64898/2026.04.29.721685

**Authors:** Ziming Chen, Chian Teng Ong, Elizabeth M. Ross

## Abstract

Bacteria require rapid adaptation under fluctuating environmental conditions. Commonly recognized global regulators enable bacteria to respond promptly to external changes, though they are either restricted to specific bacterial taxonomies or physiological statuses, suggesting that additional regulators are required for adaptation. DNA methylation is a reversible modification affecting bacterial gene regulation. However, conventional methods can only detect one DNA methylation form each round, leaving the understanding of DNA methylation in bacterial adaptation mostly unknown. This study aimed to identify genome-wide DNA methylation variation (N6-methyladenine, N4-methylcytosine, and 5-methylcytosine) in *Escherichia coli* under different culture conditions using Oxford Nanopore sequencing. DNA samples from six conditions (normal, low oxygen, low pH, high temperature, high salt, and recovery after low pH exposure) during the exponential and stationary phases were extracted. When culture conditions were compared to the normal condition, *E. coli* exhibited more differentially methylated sites during the exponential phase than in the stationary phase. During the exponential phase, the genes differentially methylated in all conditions were involved in cellular activities, such as cellular and metabolic processes. During the stationary phase, universally differentially methylated genes were associated with oxidation responses. Subsequent analysis found that although DNA methylation analysis was affected by batch effects, some genes (e.g. *rpoS*) showed consistently differential methylation across datasets. Our findings suggest that the *E. coli* DNA methylation profile was affected by growth phases and conditions, and DNA methylation profiling by Oxford Nanopore sequencing could be a potential approach for gene activity estimation in environmental samples.

**Importance:** Bacterial DNA methylation is a reversible genetic modification affecting gene regulation, enabling rapid adaptation. Three major forms in bacteria are N6-methyladenine, N4-methylcytosine, and 5-methylcytosine. Using Oxford Nanopore sequencing, we characterized genome-wide variation in these methylation types in *Escherichia coli* under six conditions (normal, low oxygen, low pH, high temperature, high salt, and recovery after low pH exposure). DNA methylation signatures in *E. coli* varied with growth conditions. Using the normal condition as a baseline, *E. coli* during the exponential phase exhibited more differentially methylated genomic loci under stress conditions compared to the stationary phase. Under stress conditions, genes with differential methylation were associated with cellular processes or oxidative responses, depending on the growth phase. Our findings reveal that the DNA methylation signature in *E. coli* was affected by growth phases and conditions, and Oxford Nanopore-based DNA methylation profiling could be a potential approach for gene activity estimation in environmental samples.

## Introduction

Since the first appearance of bacteria 3.5 billion years ago [1], the interaction between bacteria and Earth’s environment, including biotic [2] and abiotic factors [3], has been underway. Exposure to a changing environment often requires bacteria to undergo adaptation, which enables their survival and lineage preservation. Mutations, generally considered a source of organism evolution, have been widely studied for bacterial adaptation or evolution under long-term stress exposure [4–6]. Random mutations generate diverse bacterial genotypes and phenotypes to adapt or even reshape environments [7]. However, sometimes when the external stressors are removed, the mutated organisms may require several generations to revert to their original phenotypes suitable for the initial environment, due to random DNA sequence alterations. In addition, the accumulation of deleterious mutations can reduce bacterial fitness [4], which in turn damages lineage preservation. Therefore, bacteria require different adaptation strategies for short-term or temporary environmental changes.

The temporary gene modulation by regulators is an efficient approach for bacteria to adapt to fluctuating environments. The product of *rpoS* gene, RpoS, is a well-characterized global regulator in most γ-, β-, and δ-proteobacteria (e.g. *Escherichia coli*) [8]. RpoS, also known as σ^S^ or σ^38^, is a sigma factor that can initiate a general response to multiple stressors by regulating gene expression involved in metabolism and stress responses [9]. The guanosine pentaphosphate and tetraphosphate, known as (p)ppGpp, is another global regulator. (p)ppGpp exists in a wide range of bacteria and supports a stringent response to nutritional depletion and various stressors [10]. These diverse regulatory systems allow bacterial adaptation to varied environmental conditions by temporary gene regulation; however, their activities can be restricted under several circumstances. For instance, both RpoS and (p)ppGpp are degraded under fast-growing cells [11, 12]. Therefore, bacteria may require additional regulators for adaptation to fluctuating environments.

DNA methylation is another gene regulator in organisms. DNA methylation is a reversible epigenetic modification, where DNA methylases catalyse the addition of a methyl group to the DNA base. In bacteria, methylation pattern shifts within promoters and operons result in gene expression alterations [13]. Three main DNA methylation forms exist in bacteria: 5-methylcytosine (m5C), N4-methylcytosine (m4C), and N6-methyladenine (m6A) [13]. DNA methylation may enable rapid and flexible gene regulation to temporarily adapt to short-term stresses [14, 15] without reducing bacterial fitness due to random mutations. In addition to gene expression regulation, bacterial DNA methylation is also involved in genome protection and DNA repair [16].

Conventional approaches (e.g. Bisulfite sequencing) are mostly developed for detecting the m5C DNA methylation in eukaryotes [17]. Moreover, these methods can only detect one DNA methylation form at a time, thereby limiting research on DNA methylation at higher resolution, such as variation in diverse methylation forms across the genome. Currently, Oxford Nanopore Technologies (ONT) sequencing utilizes the electrical current deviations generated during sequencing to identify different nucleotides. The unique current signals from the added methyl group also enable Nanopore sequencing to identify various DNA methylation forms concurrently at the single-nucleotide level without DNA enzymatic or chemical treatment [13]. This technology overcomes the limitation of detecting a singular form and facilitates bacterial DNA methylation analyses at diverse resolutions, ranging from *de novo* DNA methylation motif identification to DNA methylation variance at the single-nucleotide level across genomes.

This study aimed to identify the genome-wide DNA methylation variation of *E. coli* under different culture conditions using Oxford Nanopore sequencing. The localized DNA methylation variance and differentially methylated genes among different culture conditions, as well as the functions of corresponding genes, were examined.

## Materials and methods

### Bacterial strains and culture conditions

The *E. coli* isolate from chicken faeces was used in this study. The isolate was cryopreserved in 25% glycerol at −80℃. Revival was performed by streaking the bacterial stock on a Luria-Bertani (LB) agar, followed by a 15-hour incubation at 37℃. A single colony from the LB agar was inoculated into a 250 ml Erlenmeyer Flask with 75 ml defined broth for cultivation in liquid media. Five LB-based culture conditions were included in this study, representing the normal culture, high temperature, high salt, low pH, and low oxygen transmission (**Table 1**).

**Table 1.** Luria-Bertani-based culture conditions.

| Characteristics | Code | Temperature | Sodium chloride | Acetic acid | Paraffin | Shaking rpm | pH |
| --- | --- | --- | --- | --- | --- | --- | --- |
| Normal | 37 | 37°C | 1.0% w/v | - | - | 200 | ~6.9 |
| Recovery from low pH | RE | 37°C | 1.0% w/v | - | - | 200 | ~6.9 |
| High salt | TNA | 37°C | 3.0% w/v | - | - | 200 | ~6.9 |
| Low pH | ACE | 37°C | 1.0% w/v | 0.5 g/l | - | 200 | ~5.5 |
| Low-oxygen transmission | PA | 37°C | 1.0% w/v | - | 75 ml overlay | 200 | ~6.9 |
| High temperature | 42 | 42°C | 1.0% w/v | - | - | 200 | ~6.9 |

For the recovery test, bacterial suspension under acetic acid was collected during the stationary phase, followed by streaking the suspension on an LB agar plate and 15-hour incubation at 37℃. A single colony from the LB agar was inoculated into 75 ml LB broth (1% NaCl), which is the normal/control condition, and cultured at 37℃ under 200 rpm.

### Growth curve construction

The *E. coli* was cultured in 75 ml defined LB broth (**Table 1**) with three biological replicates. The bacterial suspensions were collected at 1.5-hour intervals for the first 12 hours, followed by 3-hour intervals for the next 12 hours. The serially diluted bacterial suspensions were spread on the LB agar plate for the measurement of the colony-forming unit (CFU). The bacterial suspension was used for OD600 value measurement with the sterile broth as a blank. The measured OD600 values were used for 0.5-hour OD600 value imputation with the Gompertz statistical models (Figure S1). Sample collection times were determined based on the imputed OD600 values.

### Sample collection and sequencing

To generate an original dataset, new bacterial cultures were established under defined conditions (Table 1) after constructing the growth curve, with three biological replicates per condition. The bacterial suspensions were collected at specific time points derived from the imputed OD600 for each culture condition (Table S1). For the recovery culture, each replicate was derived from the corresponding replicate in the acetic acid condition. For DNA extraction, the supernatant from the suspension was removed after a 5-minute centrifugation at 16,000 x g. The pellets were collected for DNA extraction using the Puregene Kit (QIAGEN, Germany) following the manufacturer’s instructions. The extracted DNA were used for DNA library preparation with the Native Barcoding Kit 96 V14 (ONT, UK). The libraries were sequenced on the PromethION 2 Solo (ONT, UK) with a FLO-PRO114M (R10.4.1) Flow Cell. The sequencing was terminated upon reaching a coverage of 200x per sample.

To ensure a reliable validation, the validation datasets were generated independently of the original dataset at a different time period. New batches of LB broth with 3%NaCl or 0.5 g/L acetic acid were prepared and used for bacterial cultures, with three biological replicates for each condition. Bacterial suspension collection, DNA extraction, DNA library preparation, and DNA sequencing were conducted following the procedures mentioned above. The DNA library was loaded on a FLO-PRO114M (R10.4.1) Flow Cell, distinct from that used in the original dataset.

### Basecalling and alignment

ONT Dorado v0.8.0 was used to perform basecalling on the DNA sequencing data under the super accurate (SUP) model, with the DNA methylation calling function (m6A, m4C, and m5C) and adapter trimming enabled. chopper v0.10.0 [18] was used to remove reads shorter than 250 bp and mean Phred quality scores lower than 10. The filtered reads were mapped to the *E. coli* reference genome (GCF_000008865) using minimap2 v2.28 [19] with the methylation tag-retaining function enabled. SAMtools v1.13 [20] was used to extract primary mapped reads from the BAM file, followed by the subsampling to 200x coverage using Rasusa v2.0.0 [21].

The filtered data were aligned back to the *E. coli* reference genome (GCF_000008865) with minimap2 v2.28 [19] to generate a BAM file with only primary mapped reads.

### DNA methylation analysis at each segment

In the locus-based approach, DNA methylation calling was performed using ONT Modkit v0.4.5 with the BAM file and reference genome without specifying a DNA methylation motif. BEDTools v2.30.0 [22] was used to divide the reference genome into 10-kb windows. All positions in the ONT bedMethyl table generated during the DNA methylation calling were included in the analysis. Within each 10-kb window, the sum of methylated bases (N_mod_) was divided by the sum of methylated and unmethylated bases (N_valid_cov_) to calculate the regional DNA methylation fraction. An unpaired t-test was used to compare the segmental DNA methylation levels among DNA methylation forms. The segmental DNA methylation levels between growth phases were compared via a paired t-test. The Holm-Bonferroni method was applied to adjust *P*-*value* for multiple group comparisons. The coverage threshold for each genomic locus was 20x.

### De novo DNA methylation motif identification

Using the locus-based bedMethyl table and reference file, ONT Modkit v0.4.5 was applied to identify rare motifs (low occurrence and low methylation levels) with the setting developed in our previous study [23]. Since the sequencing depth was 200x, the minimum sites for considering DNA methylation motif were set as 150 (--min-sites 150) in this study.

Snappy [24] was also used for *de novo* DNA methylation motif discovery to validate the findings. In brief, DNA methylation motifs were identified using the locus-based bedMethyl file generated by ONT Modkit v0.4.5 and the reference genome file under the default settings.

### DNA methylation analysis in motifs

In the motif-based approach, DNA methylation calling was performed using ONT Modkit v0.4.5 with the BAM file and reference genome by defining a DNA methylation motif. The three most abundant DNA methylation motifs (5’-G^m6^ATC-3’, 5’-C^m5^CWGG-3’, 5’-GATCNN^m4^C-3’) from *de novo* motif identification were used in this analysis. All positions in the ONT bedMethyl table generated during the DNA methylation calling were included in the analysis. For calculating the genome-level DNA methylation level of a motif, the methylated base number (N_mod_) was divided by the sum of methylated and unmethylated bases (N_valid_cov_) of all motif loci, regardless of whether the motif loci were methylated or not. An unpaired t-test was used to compare the genome-level DNA methylation fraction among DNA methylation forms. The genome-level DNA methylation fraction between growth phases was compared via a paired t-test. The Holm-Bonferroni method was applied to adjust *the P*-*value* for multiple group comparisons. The coverage threshold for each methylation motif locus was 20x.

### Visualization of localized DNA methylation variation

The ONT bedMethyl table from the locus-based approach was used to calculate DNA methylation fraction by dividing the methylated base number (N_mod_) by the sum of methylated and unmethylated bases (N_valid_cov_) at each locus. The matrix with DNA methylation fractions was used for principal component analysis (PCA) or t-distributed Stochastic Neighbour Embedding (t-SNE) to visualize the localized DNA methylation variation among samples. The maximum iteration for t-SNE was 2000, with perplexity at 6 and theta at 0.

For each motif locus in the bedMethyl table, the DNA methylation fraction is calculated by dividing the methylated base number (N_mod_) by the sum of methylated and unmethylated bases (N_valid_cov_). The matrix with DNA methylation fractions was used for PCA or t-SNE to visualize the motif-based DNA methylation variation among samples. The maximum iteration for t-SNE was 2000, with perplexity at 4 and theta at 0.

### Differential methylation analysis

The bedMethyl tables generated from the locus-based approach were used to identify differentially methylated sites across the genome between different culture conditions or growth phases using ONT Modkit v0.4.5, with a minimum coverage of 20x. For comparison between culture conditions, the normal culture was used as a baseline, while the exponential phase was used for the growth phase comparison. Differentially methylated position was selected with the thresholds of *balanced MAP-based p-value* at 0.05 and the absolute *Cohen’s* h size effect at 0.2. After the *E. coli* GFF annotation file (GCF_000008865) conversion through AGAT v1.4.1 [25], the converted tsv file was filtered with “source_tag” as “Refseq” and “primary_tag” as “gene”. The gene body in this study was defined as the region within the open reading frame, while the promoter region was the segment 7 to 35 bp upstream of the open reading frame. The non-regulatory area in this study referred to the region outside of the gene body and promoter regions. The alignment of the differentially methylated sites to the promoter and gene body regions was performed using the filtered annotation file, followed by the identification of differentially methylated genes. Genes with one or more differentially methylated sites in corresponding gene bodies or promoters were considered differentially methylated genes in the analysis.

The GC content background of the genome could overrepresent or underrepresent the differentially methylated sites. For example, higher GC content in the genome may identify more m5C or m4C differentially methylated sites. Therefore, the differentially methylated sites were normalized by the GC content background. In brief, the m6A sites were divided by the number of AT bases, while the m4C and m5C sites were divided by the number of GC bases. An unpaired t-test was used to compare the differentially methylated sites among DNA methylation forms, while the differentially methylated sites between growth phases were compared via a paired t-test. Fitting linear models and analysis of variance (ANOVA) were also used to evaluate the effects of culture condition, growth phase transition, and various DNA methylation forms on differentially methylated sites. The Holm-Bonferroni method was applied to adjust *P-value* for multiple group comparisons.

### Functional annotation and gene enrichment analysis

SuperExactTest v1.1.0 [26] was used to evaluate whether the overlapping gene number among different groups was greater than expected, with a *P-value* threshold at 0.05. The gene enrichment was performed using ShinyGo v0.85 [27] under all available databases, with *E. coli* O157-H7 str. Sakai as the background and default settings. The False Discovery Rate (FDR) adjusted *P-value* threshold was 0.05, with 0.15 for some noted datasets due to the small gene sizes (Table S20). The top 15 pathways, ranked by fold enrichment, were used for gene function network construction by ShinyGo v0.85 [27].

### Comparison of validation and original datasets

For the validation dataset, DNA basecalling, data filtering, and DNA methylation calling were performed using the same pipeline as the original dataset. The DNA differential methylation analysis was performed using the exponential phase or acetic acid treatment group as the baseline. The comparison between the validation and original datasets was conducted through three approaches: 1) the localized DNA methylation profile; 2) the overlapping differentially methylated genes; and 3) the correlation of differentially methylated genes.

For localized DNA methylation profiling, the analysis was conducted using t-SNE and PCA based on the locus-based DNA methylation matrix, with the coverage threshold at 20x. The maximum iteration for t-SNE was 2000, with perplexity at 4 and theta at 0. For the overlapping differentially methylated gene analysis, the gene overlap significance between the origin and validation datasets was evaluated by SuperExactTest v1.1.0 [26], with the *P-value* threshold at 0.05.

For the correlation analysis, the *P-value* of each gene was used. Specifically, the *P-value* of each gene was determined as the lowest *balanced MAP-based p-value* of a differentially methylated genomic site on the corresponding gene promoter or gene body. The *P-value* was transformed using a base-10 logarithm. A fitting linear model was used to evaluate the relationship of transformed *P-value* between the origin and validation datasets.

## Results

The *E. coli* isolate from chicken faeces was used in this study. Five culture conditions based on Luria-Bertani (LB) broth were included for culturing in this study (**Table 1**). One recovery culture (RE) from the acetic acid treatment to regular LB broth was performed. The DNA methylation profiles of different culture conditions and growth phases were characterized using Oxford Nanopore sequencing (Table S1), followed by the analysis of DNA methylation levels, differentially methylated genes, and genes with condition-specific DNA methylation.

### Genome-level and regional DNA methylation showed subtle variation among treatments

A *de novo* DNA methylation motif identification was first performed using ONT Modkit v0.4.5 and Snappy [24]. Six DNA methylation motifs were confirmed among different culture conditions (5’- G^m6^<u>A</u>TC -3’, 5’-GC^m6^<u>A</u>GNNNNNTGT-3’, 5’-CCA^m6^<u>A</u>NNNNNNNNNTGG -3’, 5’-GATCNT^m4^<u>C</u>-3’, 5’-GATCNNT^m4^<u>C</u>-3’, 5’- C^m5^<u>C</u>WGG -3’), after the comparison of identified motifs and the validation with inferred motifs from the previous genome annotation [23]. To compare the genome-level DNA methylation proportions among samples, the three most abundant motifs (5’-G^m6^ATC-3’, 5’-C^m5^CWGG-3’, 5’-GATCNN^m4^C-3’) were selected. Consistent with our previous study [23], genomic DNA methylation levels were consistently lower during the exponential phase across all conditions (*P* < 0.05; Figure S2A) than in the respective stationary phase. However, significant variation was not observed between different conditions under the same growth phase.

In addition to the DNA methylation motifs, we further performed DNA methylation calling on each genomic location, which revealed DNA methylation variation at higher resolutions. Using each genomic locus, the DNA methylation level of each 10-kb genomic segment was calculated to determine the regional DNA methylation variation. Similarly, the regional DNA methylation levels were significantly lower during the exponential phase in all DNA methylation forms under different treatments (*P* < 0.001; Figure S2B). In comparison, the regional levels of most DNA methylation forms showed no differences across treatments. In summary, the DNA methylation fractions, determined at both the genome and regional levels, are lower during the exponential phase than the stationary phase, but only subtle variance was observed among different culture conditions.

### Localized DNA methylation showed variation across culture conditions and growth phases

During the adaptation, bacteria may regulate gene expression through DNA methylation on specific regulatory regions. These localized DNA methylation changes may not affect the DNA methylation pattern at the genome or segment level. In addition, because of the subtle genomic and regional DNA methylation variance among treatments, statistical methods, such as principal component analysis (PCA), utilizing the global variances may underestimate the localized DNA methylation changes (Figure S3). Therefore, t-distributed Stochastic Neighbour Embedding (t-SNE), which considers the high-dimensional localized DNA methylation differences, was performed for sample clustering. By using the locus-specific DNA methylation levels across the genome, samples were segregated into two growth phases (Figure 1A). Likewise, in the exponential phase, samples were clustered by culture condition, especially the 3% NaCl and acetic acid conditions. However, although localized variation was considered using t-SNE, samples from different treatments during the stationary phase were closely aligned. Similar patterns were seen when using DNA methylation proportions at each methylation motif position in the genome (Figure 1B). These results demonstrate that the growth phase transition is a major DNA methylation variance in *E. coli* compared to the culture conditions.

**Figure 1.**
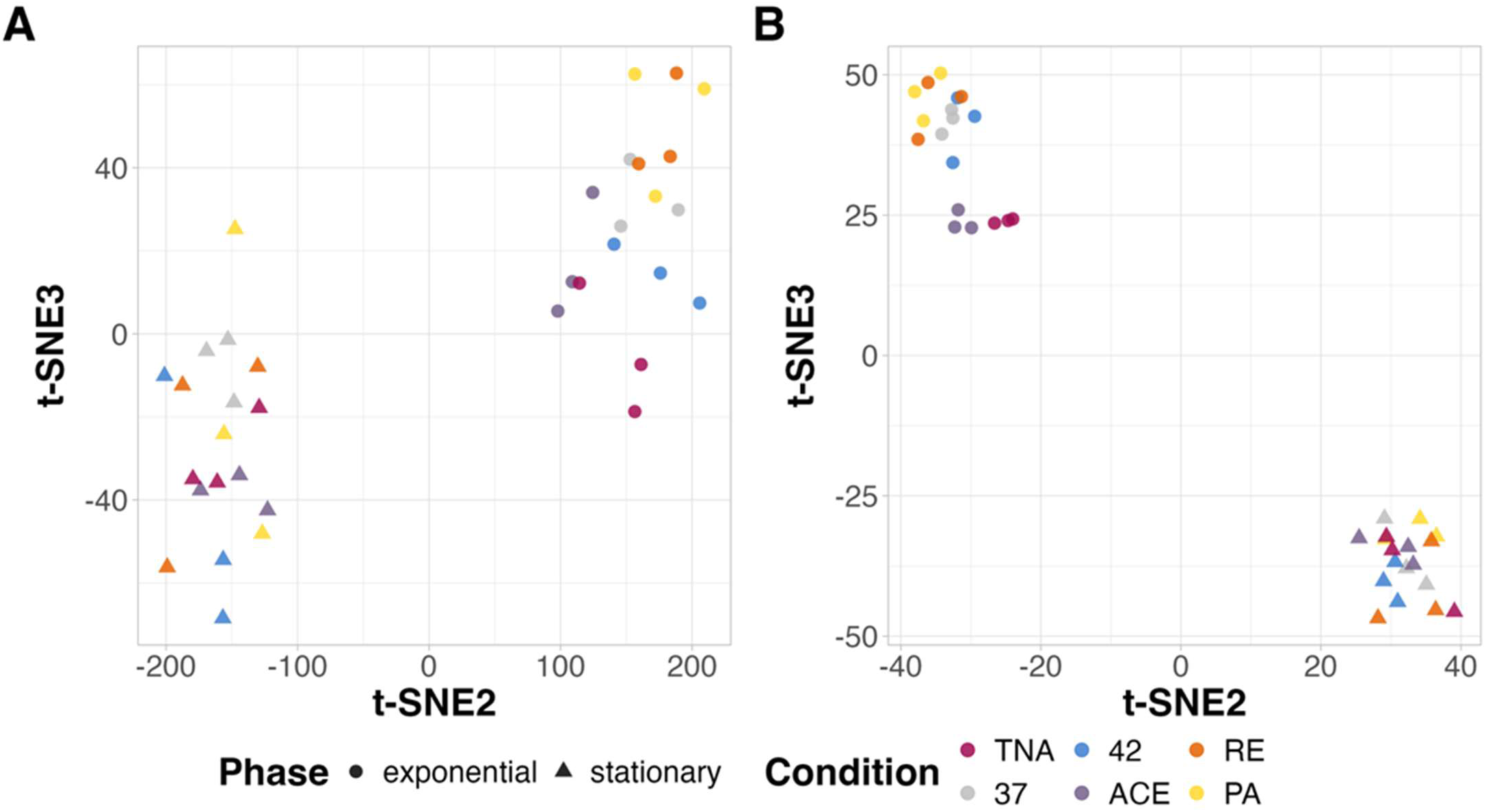
The t-distributed Stochastic Neighbour Embedding visualisation using DNA methylation levels at (A) all genomic loci and (B) motif positions. The DNA methylation level matrix for each motif position or genomic position was used for the analysis. 37: 37℃ 1%NaCl LB; 42: 42℃ 1%NaCl LB; TNA: 37℃ 3%NaCl LB; ACE: 37℃ 1%NaCl LB with 0.5 g/l acetic acid; PA: 37℃ 1%NaCl LB with paraffin overlay; RE: 37℃ 1%NaCl LB culture recovered from ACE.

### Differential methylation profiles varied across culture conditions and growth phases

Because a weak clustering pattern was observed in the stationary phase, we hypothesized that samples from different culture conditions during the stationary phase may also have fewer differentially methylated sites compared to the exponential phase. Our previous study found that using all genomic positions, regardless of the existence of a motif, revealed DNA methylation variation at higher resolutions [23]. Therefore, the differentially methylated positions were identified using all genomic positions with samples from the normal condition (37℃ 1%NaCl LB) or the exponential phase as a baseline.

Differential methylation analysis of the growth phases showed a higher number of differentially methylated m5C sites across culture conditions (*P* < 0.01; Figure 2A), compared to the other methylation forms. As expected, the number of differentially methylated sites in m6A and m4C was significantly lower in the stationary phase across culture conditions, compared to the exponential phase (*P* < 0.01; Figure 2B). In addition, the m4C showed the most differentially methylated sites across various culture conditions in both exponential and stationary phase samples (*P* < 0.001; Figure 2B). Overall, *E. coli* exhibits greater DNA methylation variation when exposed to stressors during the exponential phase.

**Figure 2.**
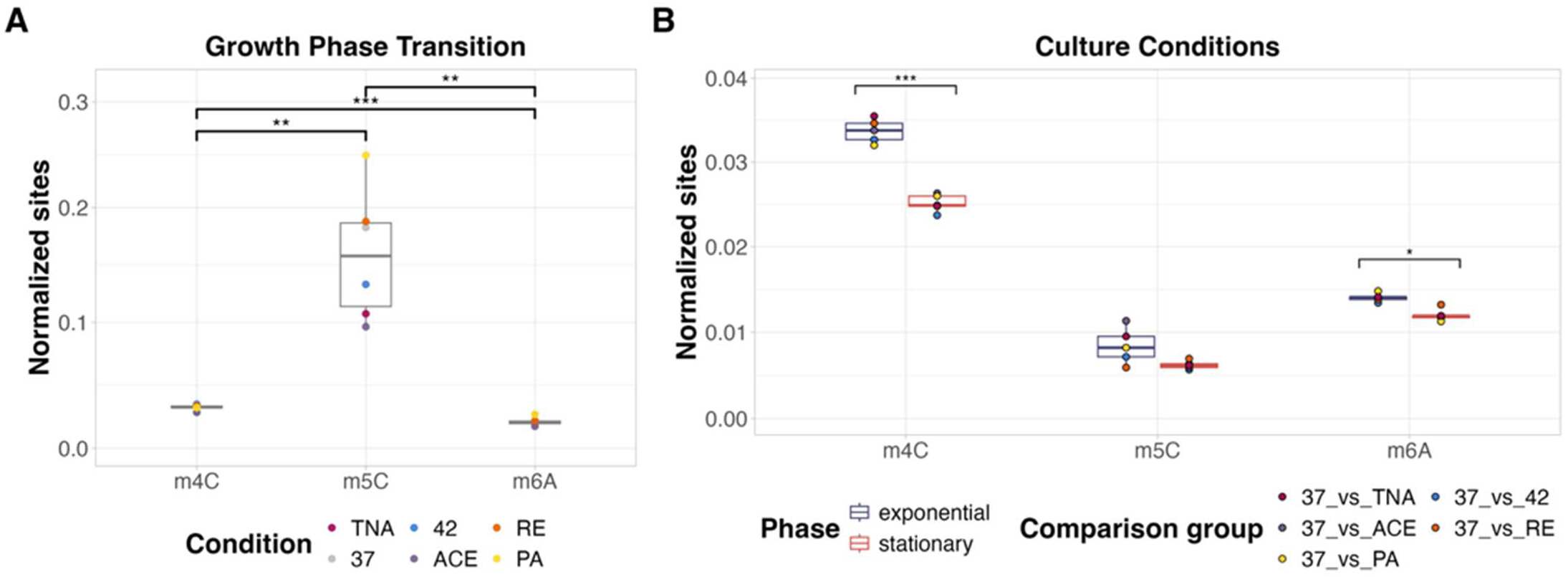
The differentially methylated sites in (A) growth phase transition and between (B) culture conditions. The identified differentially methylated sites were normalized based on GC content. A unpaired t-test was used to compare the differentially methylated sites among DNA methylation forms, while the differentially methylated sites between growth phases were compared via a paired t-test and fitting linear models. 37: 37℃ 1%NaCl LB; 42: 42℃ 1%NaCl LB; TNA: 37℃ 3%NaCl LB; ACE: 37℃ 1%NaCl LB with 0.5 g/l acetic acid; PA: 37℃ 1%NaCl LB with paraffin overlay; RE: 37℃ 1%NaCl LB culture recovered from ACE. *P*-values < 0.05: *; *P*-values < 0.01: **; *P*-values < 0.001: ***.

### Conserved differentially methylated genes were related to cellular activities

Genes with differentially methylated sites in corresponding gene bodies or promoters were considered differentially methylated genes in this analysis. The overlapping gene profiles under different growth phases and conditions were characterized. With the exponential phase samples as the baseline, 2,996 genes showed differential methylation. During the transition from the exponential to the stationary phase, 18% of genes showed consistent differential methylation across all conditions (*P* < 0.001; Figure 3A). Particularly, the *rpoS* gene, encoding the stationary-related sigma factor RpoS, was differentially methylated between growth phases across culture conditions (*P* < 0.01; Figure 4). Functions of the conserved differentially methylated genes were related to cellular activities, such as cellular anatomical entity and cellular process (*P* < 0.05; Figure 3B). In addition, several genes were associated with membrane structure, such as the plasma membrane and transmembrane.

**Figure 3.**
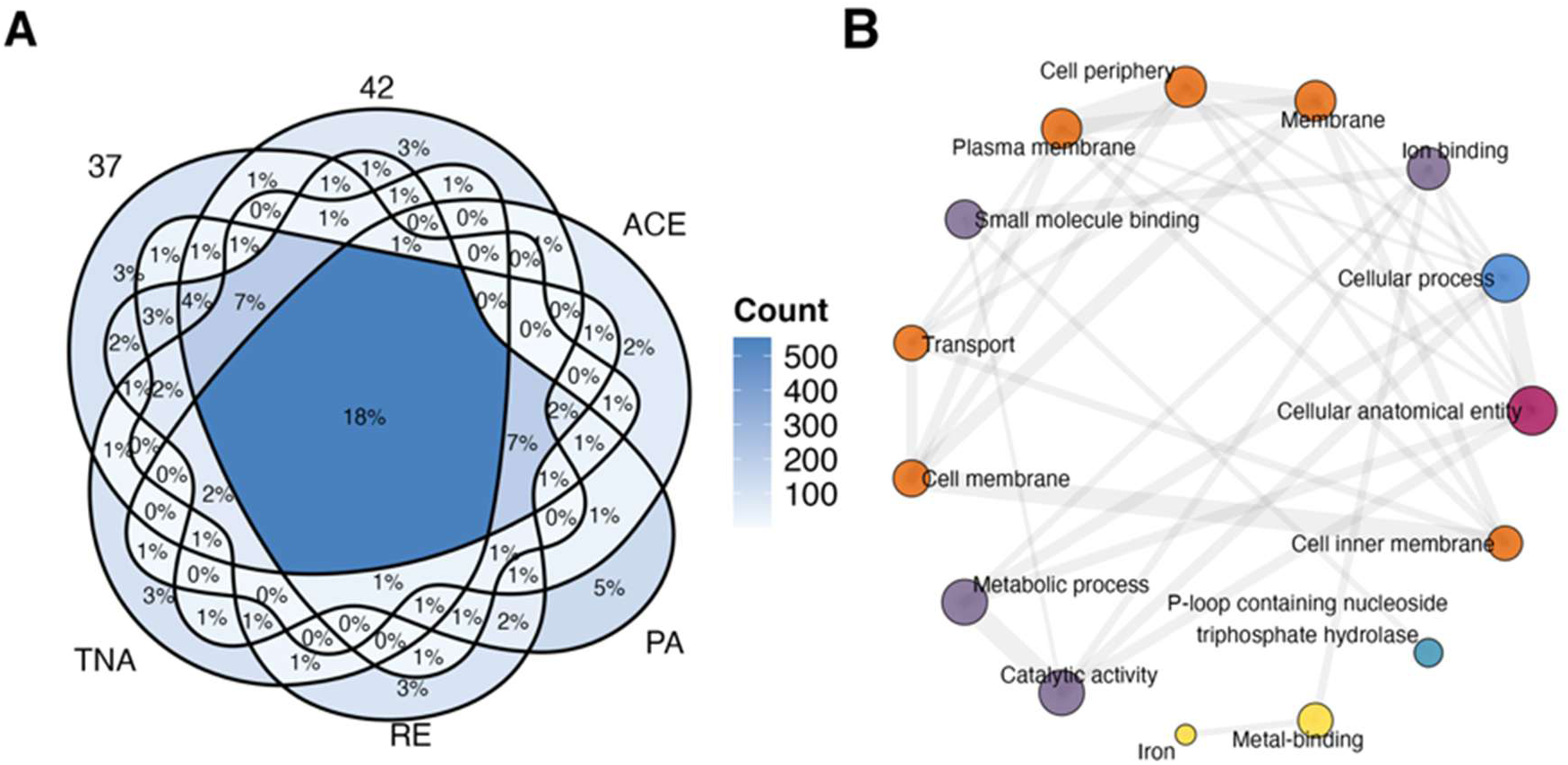
(A) The overlapping genes with differential methylation between growth phases across different conditions. (B) The function network of genes with conserved differential methylation across conditions. SuperExactTest v1.1.0 [26] performed the gene overlapping significance test. The gene enrichment analysis used the *E. coli* O157:H7 str. Sakai as a background. The top 15 pathways, ranked by fold enrichment, were used for gene function network construction. 37: 37℃ 1%NaCl LB; 42: 42℃ 1%NaCl LB; TNA: 37℃ 3%NaCl LB; ACE: 37℃ 1%NaCl LB with 0.5 g/l acetic acid; PA: 37℃ 1%NaCl LB with paraffin overlay; RE: 37℃ 1%NaCl LB culture recovered from ACE.

**Figure 4.**
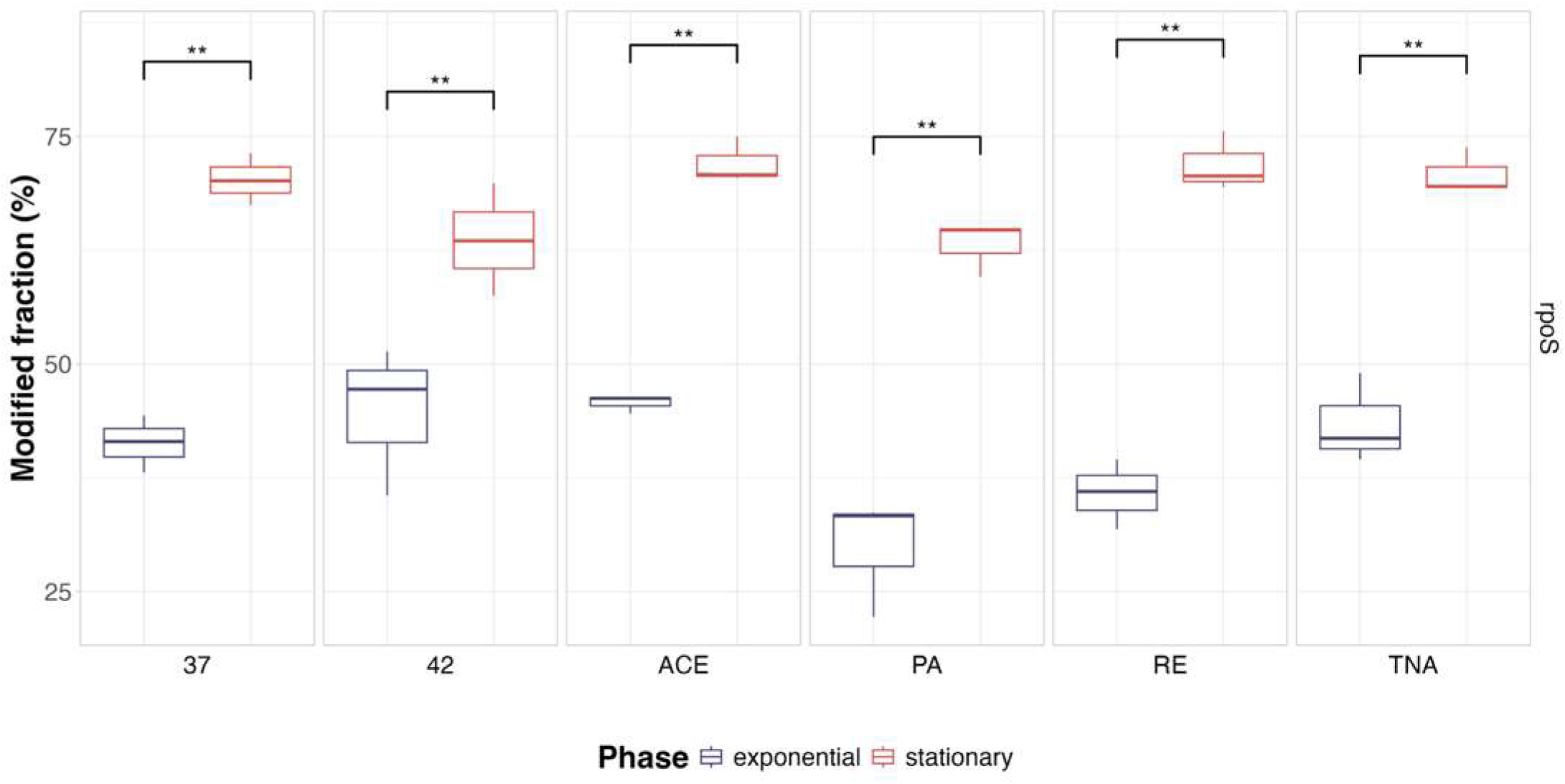
The DNA methylation levels of the *rpoS* gene at position 3,587,829 on the reverse strand of chromosome NC_002695.2. A paired t-test was used to compare the DNA methylation levels of the *rpoS* gene between the exponential and stationary phases. 37: 37℃ 1%NaCl LB; 42: 42℃ 1%NaCl LB; TNA: 37℃ 3%NaCl LB; ACE: 37℃ 1%NaCl LB with 0.5 g/l acetic acid; PA: 37℃ 1%NaCl LB with paraffin overlay; RE: 37℃ 1%NaCl LB culture recovered from ACE. *P-values* < 0.05: *; *P-values* < 0.01: **; *P-values* < 0.001: ***.

With the regular culture condition as the baseline, 2,620 and 2,498 genes showed differential methylation under exponential and stationary phases, respectively. In both the exponential and stationary phases, approximately 4% of these genes showed differential methylation across culture conditions (*P* < 0.001; Figure 5A and Figure 5C). During exponential phases, the conserved differentially methylated genes were mostly involved in cellular activities, such as cellular anatomical entity, cellular process, and metabolic process (*P* < 0.05; Figure 5B). However, in addition to cellular activities, the stationary phase had more genes associated with oxidation responses (*P* < 0.15; Figure 5D). Overall, *E. coli* exhibited greater methylation variance under stressors during the exponential phase, and conserved methylated genes were observed across growth phase transition and culture conditions.

**Figure 5.**
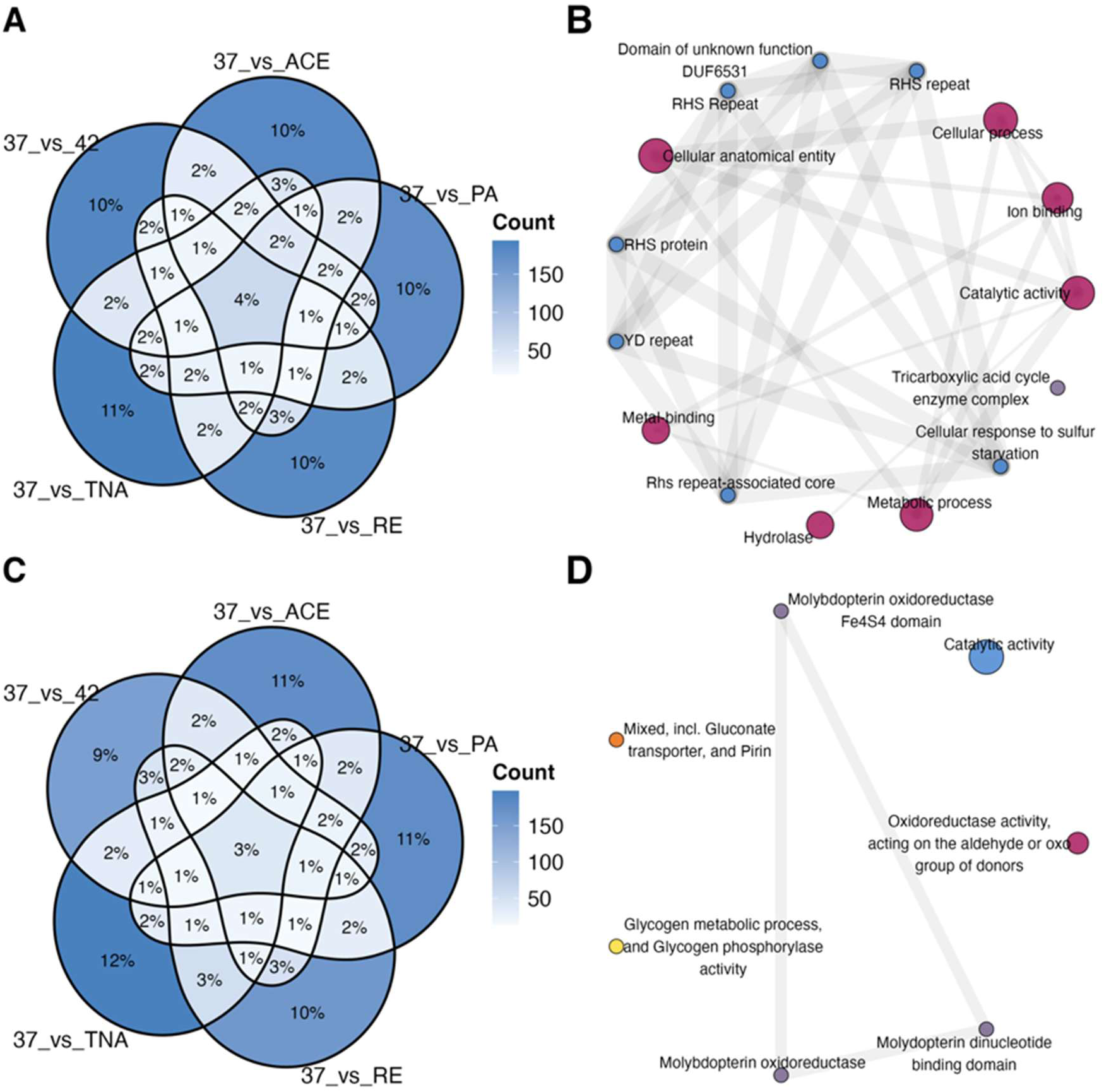
The overlapping differentially methylated gene across different conditions under the (A) exponential and (C) stationary phase. The function network of genes with conserved differential methylation across conditions under the (B) exponential and (D) stationary phase. SuperExactTest v1.1.0 [26] performed the gene overlapping significance test. The gene enrichment analysis used the *E. coli* O157:H7 str. Sakai as a background. The top 15 pathways, ranked by fold enrichment, were used for gene function network construction. 37: 37℃ 1%NaCl LB; 42: 42℃ 1%NaCl LB; TNA: 37℃ 3%NaCl LB; ACE: 37℃ 1%NaCl LB with 0.5 g/l acetic acid; PA: 37℃ 1%NaCl LB with paraffin overlay; RE: 37℃ 1%NaCl LB culture recovered from ACE.

### Differentially methylated genes were specific to culture conditions

In addition to the general stress response, the adaptation to an environment may also require additional regulators [9], such as DNA methylation. Therefore, the uniquely differentially methylated genes under each growth culture and their functions were identified. Around 50% of the genes, such as *cusF* and *fepD*, were culture-condition specific in both growth phases, with the normal condition as the baseline (Figure 5A and Figure 5C; Table S13). In addition, samples restored to the normal condition (RE) after the acetic acid challenge (ACE) demonstrated a distinct methylated gene profile (Figure 5A and Figure 5C). However, 39.60% the differentially methylated genes in the recovery group (RE) were also present in the acetic acid group (ACE) during the exponential phase, and 36.43% during the stationary phase. Functional annotation analysis revealed that they are mostly related to cellular anatomical entity or cellular process (*P* < 0.15; Figure S7 and Figure S8; Table S18).

### Validation of the growth-related DNA methylation

To examine the batch effect on DNA methylation profiling, validation datasets were generated independently of the original dataset at a different time period. The validation dataset consisted of two growth settings: 3%NaCl and 0.5 g/L acetic acid. The genome-wide DNA methylation variance between the origin and validation datasets was compared. Both t-SNE and PCA revealed a clear separation between the origin and validation datasets (Figure S9); however, growth transition still accounts for a large DNA methylation variance in both datasets. These results demonstrate that samples from different batches can introduce variation in DNA methylation profiling.

The differential methylation analysis was performed using samples from the same batch, with the exponential phase or acetic acid treatment (ACE) group as the baseline. Despite batch effects, some genes were universally differentially methylated (*P* ≤ 0.05) across both datasets, with growth phase transition exhibiting the greatest consistency (*P* < 0.001; Figure 6A and Figure 6B). For instance, the *rpoS* gene, encoding the RpoS sigma factor, consistently showed differential methylation on the reverse strand at position 3,587,829 of chromosome NC_002695.2 across all samples in both datasets (*P* < 0.001; Table S21). Although the number of mutual genes with differential methylation was lower when comparing the acetic acid (ACE) and high salt (TNA) treatments, the valuation and origin are still significantly correlated (*P* < 0.001; Figure 6C and Figure 6D). These findings show that batch effects influence DNA methylation analysis, yet positive correlations remain between batches.

**Figure 6.**
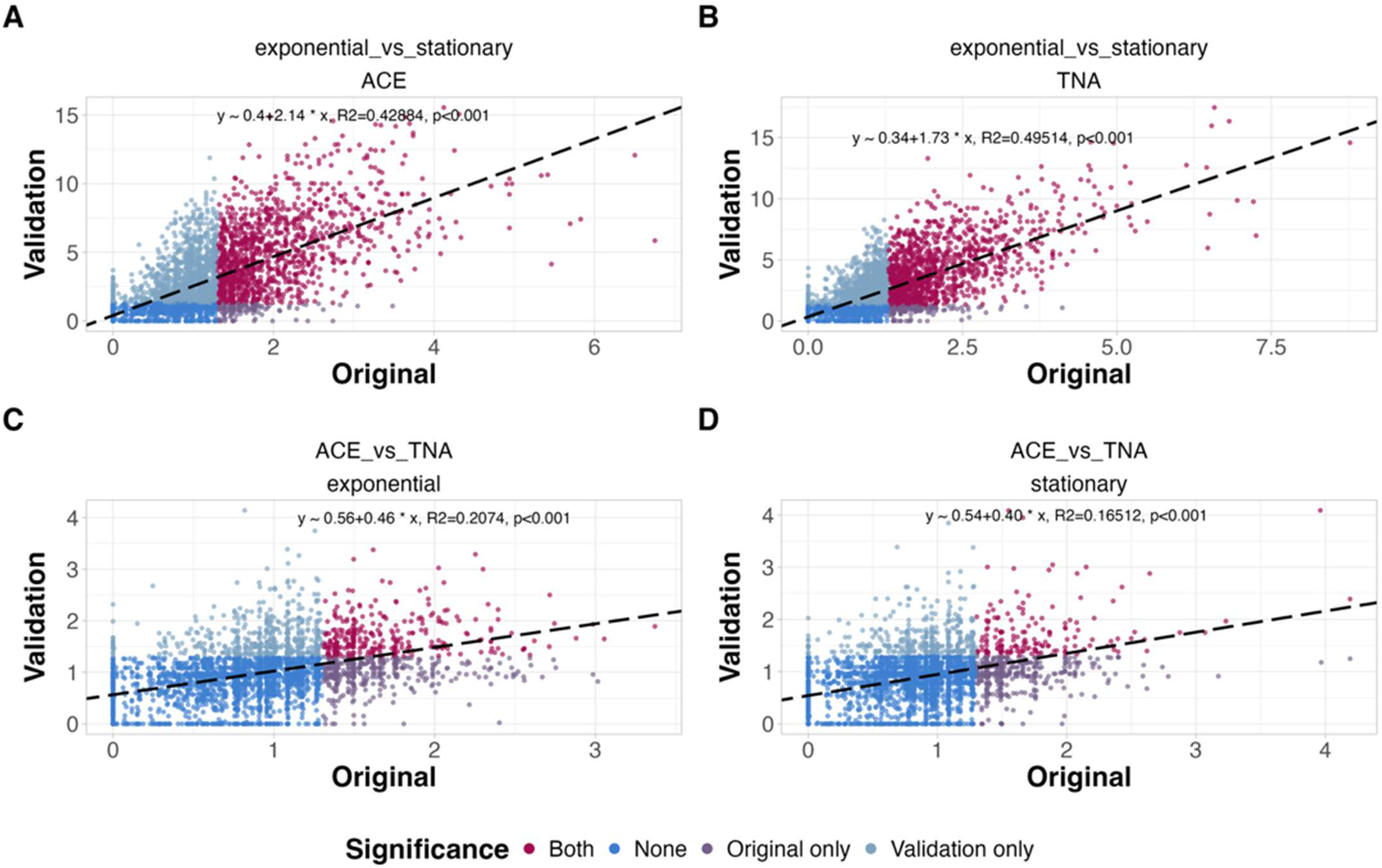
The correlation of differentially methylated gene *P-values* between the origin and validation datasets for growth-phase (A, B) and culture-condition (C, D) comparisons. The *P-value* for each gene was determined as the lowest *balanced MAP-based p-value* for a differentially methylated genomic site on the corresponding gene promoter or gene body. The *P-values* were transformed by base-10 logarithm. A fitting linear model was used to estimate the correlation between datasets. TNA: 37℃ 3%NaCl LB; ACE: 37℃ 1%NaCl LB with 0.5 g/l acetic acid.

## Discussion

This study investigated the genome-scale DNA methylation variation (m6A, m4C, and m5C) in *E. coli* under six culture conditions and two growth phases. We found that in *E. coli*, DNA methylation variation between different culture conditions was smaller than DNA methylation variation between growth phases. The functions of genes with universal differential methylation in response to growth conditions differed from those under growth phase transition. During the exponential phase, the universal differentially methylated genes were involved in various cellular activities, such as cellular anatomical entity, cellular process, and metabolic process. During the stationary phase, the universal differentially methylated genes were associated with the oxidation response. Multiple differentially methylated genes, such as *rpoS* and *cusF*, were specific to growth phases or culture conditions for general stress response and iron/ion transport. Batch effects were seen between the original and validation datasets; however, some genes (e.g. *ropS*) consistently showed differential methylation across datasets. Our findings suggest that the *E. coli* DNA methylation profile varies across growth phases and culture conditions, and DNA methylation profiling by Oxford Nanopore sequencing could be an efficient approach for gene activity in bacteria, particularly *E. coli*.

DNA methylation variance in *E. coli* across culture conditions was lower during the stationary phase, compared to the exponential phase. Similar to a previous study using *E. coli* transcriptomes [28], our study found that the stationary phase exhibited fewer differentially methylated genomic loci and genes than the exponential phase. This can be caused by the increased expression of *rpoS* gene (encoding the RpoS protein) during the stationary phase [29]; however, the *rpoS* gene is usually repressed during the exponential phase [12]. The RpoS protein can interact with RNA polymerase to form a holoenzyme called Eσ^S^ [30]. RpoS can direct RNA polymerase to the specific promoter sequence 5’-TCTATACTTAA-3’ [9], enabling efficient gene transcription for stress response under the stationary phase [30, 31]. Multiple stress response genes, such as *dps* and *osmC*, can be regulated by the interaction between the Eσ^S^ and the promoter sequence [32] to initiate a general stress response, supporting a broad resistance to various stressors during the stationary phase [12, 33]. Consistent with a previous study [34], the m5C levels at the *rpoS* gene body were consistently higher in the stationary phase than in the exponential phase across all culture conditions in our study, potentially reflecting its differential expression (although we did not test this specifically). Therefore, the general adaptation strategy by RpoS in *E. coli* may result in the lower variance in transcriptomic or methylation profiles across culture conditions during the stationary phase.

Similar to previous findings on *E. coli* transcriptomes or metabolomics [28, 35], the growth phase transition contributed the most DNA methylation variance in our study, regardless of the culture condition. These can be caused by the systematically lower DNA methylation levels during the exponential phase compared to the stationary phase, as was found in our study and other previous studies [23, 34]. In addition, the notable variance in DNA methylation may indicate its regulatory role in systematic cellular regulation during the growth phase transition. Our analysis found that the growth phase comparison yielded 376 more differentially methylated genes than the culture condition comparison. Consistent with previous studies [9, 35], these differentially methylated genes were mostly related to the cell membrane and metabolism, allowing energy conservation during the general stress response during the stationary phase [9, 35]. In real scenarios, microbes are predominantly in the stationary phase in nature. The transition to exponential growth generally occurs under specific conditions, such as acute infections. Therefore, the substantial DNA methylation variation during the growth phase transition may mask signals from other environmental factors when mixed communities are studied. While we can control this during the analysis using pure microbial culture, it may not be possible in microbiome studies.

Similar to previous studies using transcriptomic data [36–40], multiple genes (e.g. *cusF*, *fepD* and cheW) showed unique differential methylation in a culture condition, compared to the normal condition. The *cusF* gene, encoding the CusF protein, was differentially methylated in the paraffin overlay culture (PA) under the exponential phase. The increased *cusF* expression was considered an alternative way for regulating Cu ion concentration due to impaired CueO during anaerobic growth [40]. The *fepD* gene (Ferric enterobactin transport system permease protein, FepD) showed differential methylation under the acetic acid treatment (ACE) during the stationary phase, compared to the normal culture. The regulation of *fepD* was potentially required for bacterial adaptation to acidic environments with limited manganese and iron [38], as observed in the stationary phase here, where most of the nutrients were consumed. These findings suggest that DNA methylation patterns of *E. coli* vary under different culture conditions.

However, some of the representative genes, such as *fabA* and *fabB*, under the moderate acidic treatment (ACE) and the exponential phase, did not show differential methylation in our study. This may result from the bacterial gene regulatory diversity being independent of DNA methylation. For example, the acid-tolerance response (ATR) system protects *E. coli* under moderate acid stress (pH 4.5-5.8) [41]. The ATR system relies on the CpxRA system, consisting of a cytoplasmic response regulator CpxR and a sensor histidine kinase CpxA. The CpxRA system is activated by moderate acid, and the phosphorylated CpxR can upregulate the expression of *fabA* or *fabB* by binding to their promoter regions [41]. The overexpression of *fabA* or *fabB* enhances the unsaturated fatty acid (UFA) biosynthesis, thereby increasing membrane UFA content and restoring the growth ability of *E. coli* during the exponential phase [41]. Therefore, DNA methylation may not reveal the bacterial gene regulation complexity; however, its ability to identify culture condition-specific differentially methylated genes still enables the general estimation of bacterial physiological status.

Our findings suggest that DNA methylation profiling by Oxford Nanopore sequencing can be an alternative for microbial activity estimation in environmental samples. Generally, metagenomics is used to quantify the relative abundance of microbial communities in an environment, yet it fails to measure gene activity. Therefore, other omics data, including metatranscriptomics [42] and metaproteomics [43], have been incorporated to evaluate microbial activity across various environments. These approaches directly measure the abundance of gene products or metabolites, providing a relatively accurate evaluation of microbial activities; however, their implementation is complicated by several technical challenges. For instance, the mRNA instability and the absence of poly(A) tails in prokaryotic mRNA increase the difficulty in sample preparation for metatranscriptomics [42]. Likewise, the high host-derived protein abundance also complicates and prolongs sample preparation for metaproteomics [43]. Encouragingly, Oxford Nanopore sequencing can provide two types of omics data in a single run: genomics and epigenomics, making the assay very cost-effective compared to other methods. Our findings also show that the DNA methylation profiles of several *E. coli* genes vary under different growth phases and culture conditions. Although only one bacterial species was used in our study, Nanopore DNA methylation profiling provides a potential approach for gene activity estimation, which could potentially be applied to microbiome samples.

The post-acid *E. coli* (RE), where the cells were cultured in an acid solution and then inoculated into a normal LB broth, exhibited a distinct DNA methylation profile. Notably, a proportion of differentially methylated genes was shared with samples under acidic (ACE) stress, especially during the stationary phase. The distinct DNA methylation profiles observed in the recovery group may reflect cellular heterogeneity during recovery process. This could explain, for example, why a previous study found that only around 50% of the *Pseudomonas aeruginosa* isolates evolved *in vivo* reverted to the ancestor morphology after a 38-generation recovery [44]. These findings suggest that single-cell DNA methylation analysis in bacteria may allow methylation tracking at higher resolution and enable the measurement of the time required for DNA methylation recovery after stress challenges. On the other hand, these findings indicate that bacteria may encode the “memory” of prior environmental conditions using DNA methylation [45]. Other mechanisms have been proposed for the bacterial memory system, such as iron-based [46] and cAMP-TFP-based [47]. These memories can be passed on for several generations after exposure to a different environment [45–47]. Therefore, the bacterial memory system may enable rapid and energy-conserved bacterial adaptation when confronting the same stressor.

Similar to a previous study using methylation data by Illumina [48], DNA methylation profiling by Oxford Nanopore sequencing was also affected by batch effects in our study. Batch effects refer to variance resulting from technical variations during the experiment, such as sample preparation procedures, experiment equipment, and operators [49]. Therefore, differences between the origin and validation datasets in our study likely resulted from the use of different batches of culture media and flow cells. In addition to DNA methylation, the batch effect also introduced significant variance to other omics studies, including transcriptomics [50] and genomics [51]. These findings indicate that incorporating data from different sources without batch effect correction can affect data analysis and draw biased conclusions. Appropriate analysis methods are necessary for a study using datasets from different batches, such as careful study design, the inclusion of an internal control group, and batch effect correction.

## Conclusion

This study characterized *E.coli* DNA methylation profiles, including localized DNA methylation and differentially methylated genes, under different growth phases and culture conditions. The growth phase transition introduced more DNA methylation variance compared to the culture conditions. The differentially methylated genes reflected the *E. coli* cellular response during growth phase transition and under different stressors, although batch effects were observed. Our findings suggest that the *E. coli* DNA methylation profile is affected by growth phases and culture conditions, and DNA methylation profiling by Oxford Nanopore sequencing could be a potential approach for gene activity estimation in environmental samples.

## Availability of data and materials

The raw sequence data of this study were archived under the NCBI BioProject: PRJNA1450215. Sample accession numbers are listed in Table S24.

The POD5 data for this study are available on UQ eSpace: https://doi.org/10.48610/1ae1346

Coding scripts were available on the GitHub repository: https://github.com/SimonChen1997/DNA_methylation_stress_response

## Acknowledgements

We would like to thank the colleagues at the Centre for Animal Science (Queensland Alliance for Agriculture and Food Innovation, University of Queensland) for providing valuable feedback for this study. We acknowledge the support from the Australian Government Research Training Program Scholarship.

## Contributions

Conception and design were by Z.C, C.T.O, and E.M.R. Sample collection and experiments were by Z.C and C.T.O. Data analysis and visualization were by Z.C, C.T.O, and E.M.R. Draft writing was by Z.C. Draft reviewing was by Z.C, C.T.O, and E.M.R.

